# Hetnet connectivity search provides rapid insights into how two biomedical entities are related

**DOI:** 10.1101/2023.01.05.522941

**Authors:** Daniel S. Himmelstein, Michael Zietz, Vincent Rubinetti, Kyle Kloster, Benjamin J. Heil, Faisal Alquaddoomi, Dongbo Hu, David N. Nicholson, Yun Hao, Blair D. Sullivan, Michael W. Nagle, Casey S. Greene

## Abstract

Hetnets, short for “heterogeneous networks”, contain multiple node and relationship types and offer a way to encode biomedical knowledge. One such example, Hetionet connects 11 types of nodes — including genes, diseases, drugs, pathways, and anatomical structures — with over 2 million edges of 24 types. Previous work has demonstrated that supervised machine learning methods applied to such networks can identify drug repurposing opportunities. However, a training set of known relationships does not exist for many types of node pairs, even when it would be useful to examine how nodes of those types are meaningfully connected. For example, users may be curious not only how metformin is related to breast cancer, but also how the *GJA1* gene might be involved in insomnia. We developed a new procedure, termed hetnet connectivity search, that proposes important paths between any two nodes without requiring a supervised gold standard. The algorithm behind connectivity search identifies types of paths that occur more frequently than would be expected by chance (based on node degree alone). We find that predictions are broadly similar to those from previously described supervised approaches for certain node type pairs. Scoring of individual paths is based on the most specific paths of a given type. Several optimizations were required to precompute significant instances of node connectivity at the scale of large knowledge graphs. We implemented the method on Hetionet and provide an online interface at https://het.io/search. We provide an open source implementation of these methods in our new Python package named [hetmatpy](https://github.com/hetio/hetmatpy).

## Introduction

A *network* (also known as a graph) is a conceptual representation of a group of entities — called *nodes —* and the relationships between them — called *edges*. Typically, a network has only one type of node and one type of edge. However, in many cases, it is necessary to be able to distinguish between different types of entities and relationships.

A *hetnet* (short for **het**erogeneous information **net**work [1]) is a network where nodes and edges have type. The ability to differentiate between different types of entities and relationships allows a hetnet to describe more complex data accurately. Hetnets are particularly useful in biomedicine, where it is important to capture the conceptual distinctions between various entities, such as genes and diseases, and linkages, such as upregulation and binding.

The types of nodes and edges in a hetnet are defined by a schema, referred to as a metagraph. The metagraph consists of metanodes (types of nodes) and metaedges (types of edges). Note that the prefix *meta* refers to the type (e.g. compound), as opposed to a specific node/edge/path itself (e.g. acetaminophen).

One such network is Hetionet, which provides a foundation for building hetnet applications. It unifies data from several different, disparate sources into a single, comprehensive, accessible, common-format network. The database is publicly accessible without login at https://neo4j.het.io. The Neo4j graph database enables querying Hetionet using the Cypher language, which was designed to interact with networks where nodes and edges have both types and properties.

One such application, Project Rephetio, focused on drug repurposing [2]. The authors [predicted](https://het.io/repurpose/) the probability of drug efficacy for 209,168 compound–disease pairs. A supervised machine learning approach identified types of paths that occur more or less frequently between known treatments than non-treatments (Figure 1B). To train the model, the authors created [PharmacotherapyDB](https://doi.org/10.6084/m9.figshare.3103054), a physician-curated catalog of 755 disease-modifying treatments [3].

**Figure 1:**
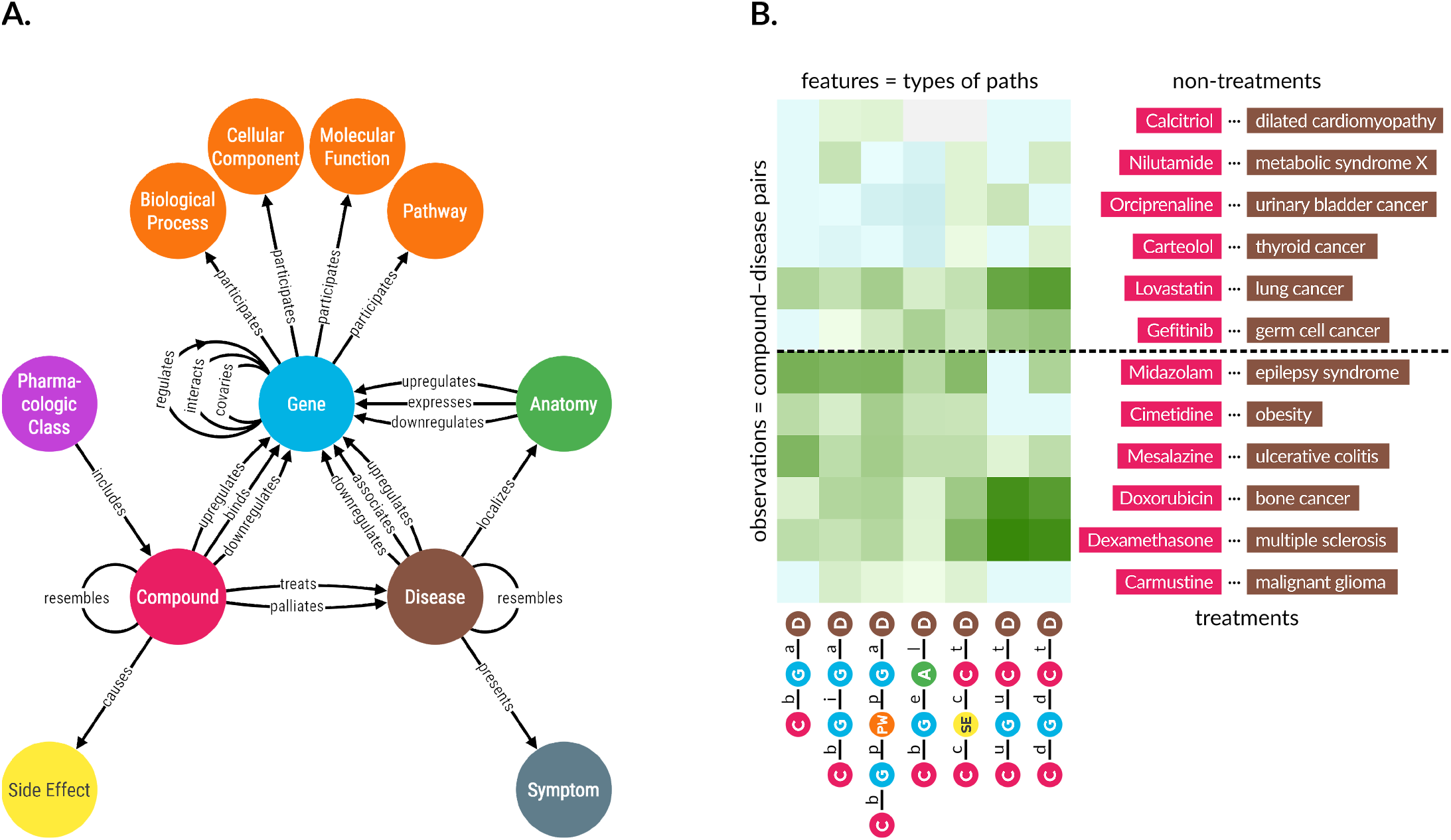
**A. Hetionet v1.0 metagraph.** The types of nodes and edges in Hetionet. **B. Supervised machine learning approach from Project Rephetio.** This figure visualizes the feature matrix used by Project Rephetio to make supervised predictions. Each row represents a compound–disease pair. The top half of rows correspond to known treatments (i.e. positives), while the bottom half correspond to non-treatments (i.e. negatives under a *closed-world assumption*, not known to be treatments in PharmacotherapyDB). Here, an equal number of treatments and non-treatments are shown, but in reality the problem is heavily imbalanced. Project Rephetio scaled models to assume a positive prevalence of 0.36% [2,4]. Each column represents a metapath, labeled with its abbreviation. Feature values are DWPCs (transformed and standardized), which assess the connectivity along the specified metapath between the specific compound and disease. Green colored values indicate above-average connectivity, whereas blue values indicate below average connectivity. In general, positives have greater connectivity for the selected metapaths than negatives. Rephetio used a logistic regression model to learn the effect of each type of connectivity (feature) on the likelihood that a compound treats a disease. The model predicts whether a compound-disease pair is a treatment based on its features, but requires supervision in the form of known treatments.

Project Rephetio successfully predicted treatments, including those under investigation by clinical trail. However, two challenges limit the applicability of Rephetio. First, Rephetio required known labels (i.e. treatment status) to train a model. Hence, the approach cannot be applied to domains where training labels do not exist. Second, the DWPC metric used to assess connectivity is sensitive to node degree. The Rephetio approach was incapable of detecting whether a high DWPC score indicated meaningful connectivity above the level expected by the background network degrees. Here we develop Hetnet connectivity search, which defines a null distribution for DWPCs that accounts for degree and enables detecting meaningful hetnet connectivity without training labels.

Existing research into methods for determining whether two nodes are related primarily focuses on homogeneous networks (without type). Early approaches detected related nodes by measuring neighborhood overlap or path similarity between two nodes [5,6]. These approaches predicted node relatedness with success. However, they are challenging to scale as a network grows in size or semantic richness (i.e. type) [5].

More recently, focus has shifted to graph embeddings to determine if two nodes are related, specifically in the context of knowledge graphs, which are often semantically rich and include type [7,8,9,10,11]. These types of methods involve mapping nodes and sometimes edges to dense vectors via neural network models [12,13,14], matrix factorization [15,16], or translational distance models [17]. Bioteque creates node embeddings from the bipartite network of DWPCs for a given metapath [18]. Once these dense vectors have been produced, quantitative scores that measure node relatedness can be generated via a machine learning model [8,19,20] or by selected similarity metrics [7,9,21,22,23]. These approaches have been quite successful in determining node relatedness. Yet, they only state *whether* two nodes are related and fail to explain *why* two nodes are related.

Explaining why two nodes are related is a non-trivial task because approaches must output more than a simple similarity score. The first group of approaches output a list of ranked paths that are most relevant between two nodes [24,25,26]. For example, the FAIRY framework explains for why items appear on a user’s social media feed based on a network of users and content classes (e.g. categories, user posts, songs) [25]. ESPRESSO explains how two sets of nodes are related by returning subgraphs [27]. Other approaches such as MetaExp return important metapaths rather than paths, but require some form of supervision [28,29]. Hetnet connectivity search explains how two nodes are related in an unsupervised manner that captures the semantic richness of edge type and returns results in the form of both metapaths and paths. Our open source implementation, including for a query and visualization webserver, was designed with scalability and responsiveness in mind allowing in-browser exploration.

## Results

Completing hetnet connectivity search involved advances on three fronts. We implemented new software for efficient matrix-based operations on hetnets. We developed strategies to efficiently calculate the desired connectivity score under the null. We designed and developed a web interface for easy access to the connectivity search approach.

### Hetmatpy Package

We created the hetmatpy Python package, available on [GitHub](https://github.com/hetio/hetmatpy) and [PyPl](https://pypi.org/project/hetmatpy/) under the permissive BSD-2-Clause Plus Patent License. This package provides matrix-based utilities for hetnets. Each metaedge is represented by a distinct adjacency matrix, which can be either a dense Numpy array or sparse SciPy matrix (see HetMat architecture). Adjacency matrices are stored on disk and loaded in a lazy manner to help scale the software to hetnets that are too large to fit entirely in memory.

The primary focus of the package is to provide compute optimized and memory efficient implementations of path counting algorithms. Specifically, the package supports computing *degree-weighted* path counts (DWPCs), which can be done efficiently using matrix multiplication but require complex adjustments to avoid counting paths with duplicate nodes (i.e. to filter walks that are not paths, see <u>DWPC matrix multiplication algorithms</u>). The package can reuse existing path count computations that span segments of a longer metapath. The package also supports generating null distributions for DWPCs derived from permuted networks, see Degree-grouping of node pairs. Since this approach generates too many permuted DWPC values to store on disk, our implementation retains summary statistics for each degree-group that allow computation of a <u>Gamma-hurdle distribution</u> from which null DWPC *p*-values can be generated.

### DWPC null distribution

To assess connectivity between a source and target node, we use the DWPC (degree-weighted path count) metric. The DWPC is similar to path count (number of paths between the source and target node along a given metapath), except that it downweights paths through high degree nodes. Rather than using the raw DWPC for a source-metapath-target combination, we transform the DWPC across all source-target node pairs for a metapath to yield a distribution that is more compact and amenable to modeling [30].

Previously, we had no technique for detecting whether a DWPC value was exceptional. One possibility is to evaluate the DWPCs for all pairs of nodes and select the top scores (e.g. the top 5% of DWPCs). Another possibility is to pick a transformed DWPC score as a cutoff. The shortcomings of these methods are twofold. First, neither the percentile nor absolute value of a DWPC has inherent meaning. To select transformed DWPCs greater than 3.5, or alternatively the top 1% of DWPCs, is arbitrary. Second, comparing DWPCs between node pairs fails to account for the situation where high-degree node pairs are likely to score higher, solely on due to their degree (5).

To address these shortcomings, we developed a method to compute the right-tail *p*-value of a DWPC. *p*-values have a broadly understood interpretation — in our case, the probability that a DWPC equal to or greater than the observed DWPC could occur under a null model. Our null model is based on DWPCs generated from permuted networks, where edges have been randomized in a degree-preserving manner (see <u>Permuted hetnets</u>).

By tailoring the null distribution for a DWPC to the degree of its source and target node (see Degree-grouping of node pairs), we account for degree effects when determining the significance of a DWPC.

To improve the accuracy of DWPC *p*-values, we use fìt a <u>gamma-hurdle distribution</u> to the null DWPCs. In rare cases, there are insufficient nonzero null DWPCs to fìt the gamma portion of the null distribution. In these cases, we fallback to an empirical calculation as described in <u>Empirical DWPC p-values</u>.

### Enriched metapaths

For each of the 2,205 metapaths in Hetionet v1.0 with length **≤** 3, we computed DWPCs for all node pairs and their corresponding null distributions, see DWPC and null distribution computation. We store the most significant DWPCs as described in <u>Prioritizing enriched metapaths for database storage</u>, which appear as the “precomputed” rows in the webapp metapath table (Figure 4B & 2). DWPCs that are not retained by the database can be regenerated on the fly. This design allows us to immediately provide users with the metapaths that are most enriched between two query nodes, while still allowing on-demand access to the full metrics for all metapaths with length **≤** 3.

**Figure 2:**
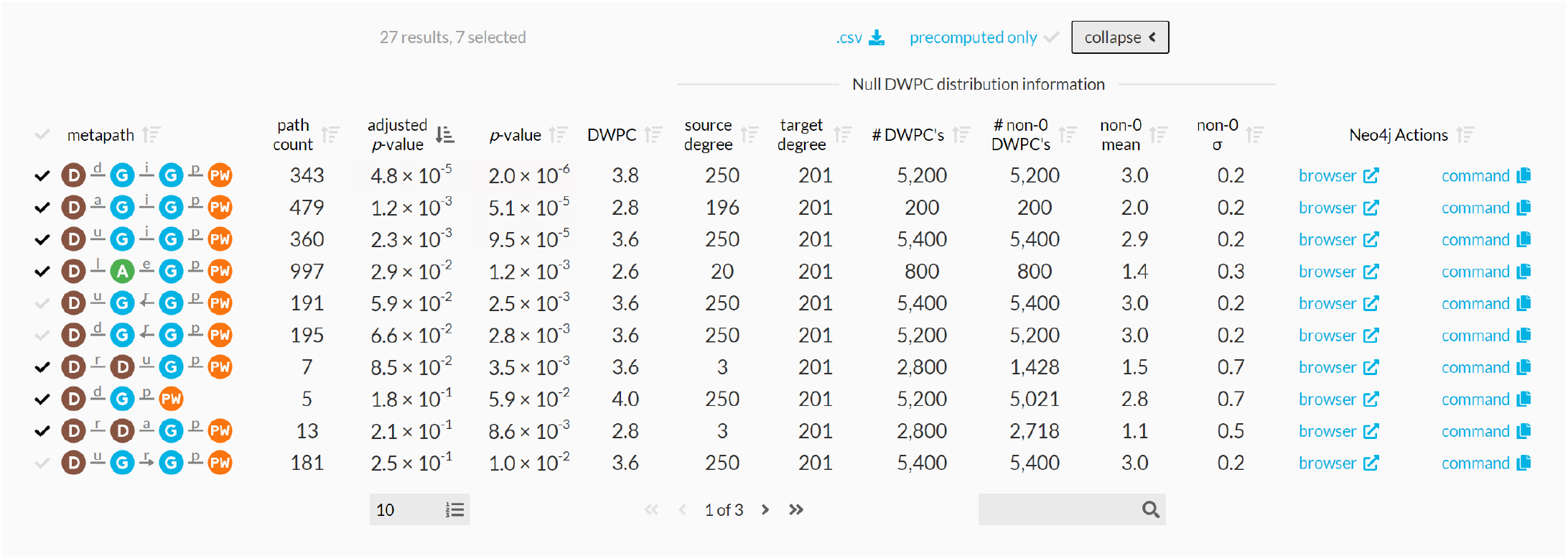
Expanded metapath details from the connectivity search webapp. This is the expanded view of the metapath table in 4B.

**Figure 3:**
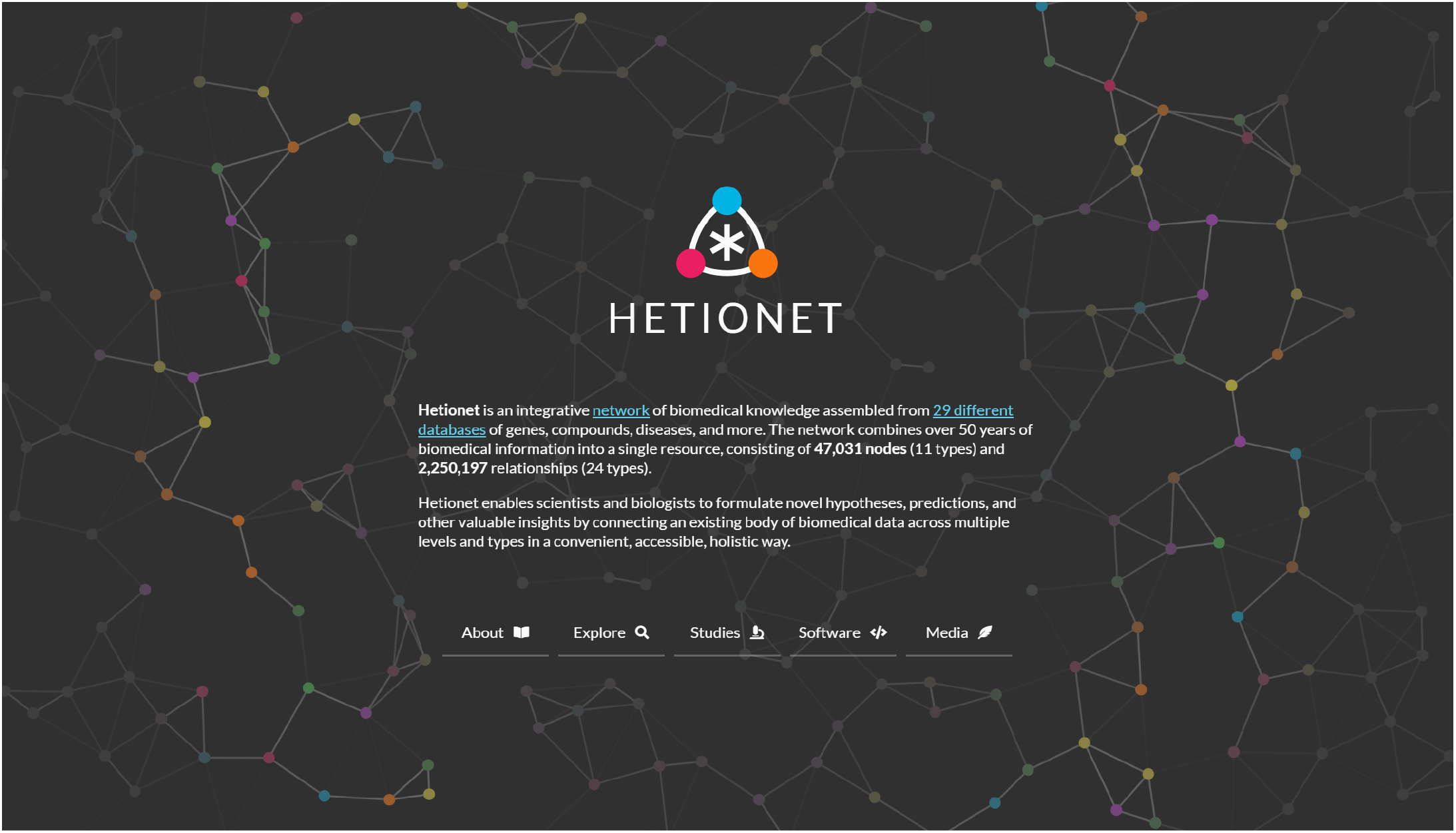
Homepage of the Hetio website. Provides a succinct overview of what Hetionet consists of and what its purpose is.

**Figure 4:**
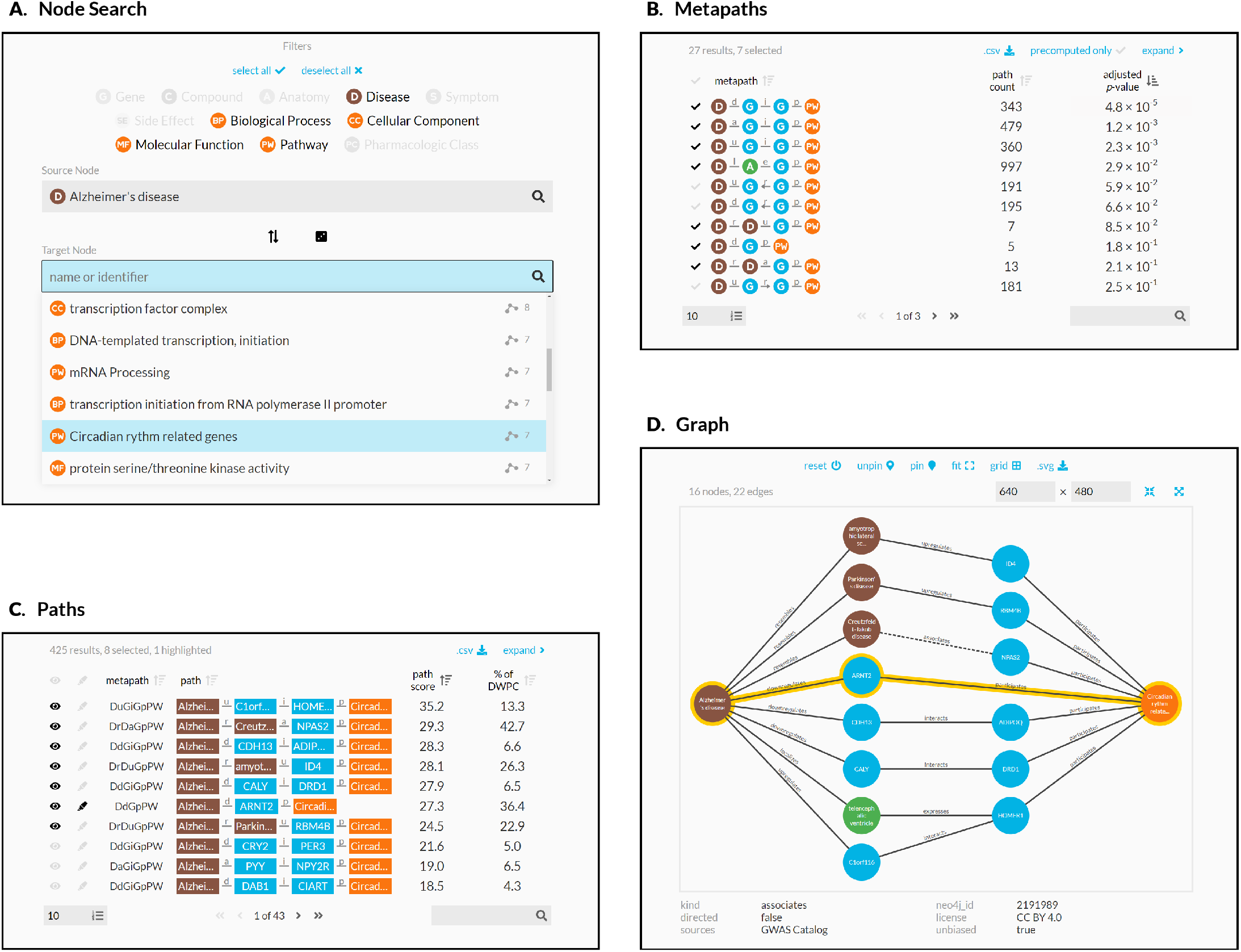
Using the connectivity search webapp to explore the pathophysiology of Alzheimer’s disease. This figure shows an example user workflow for https://het.io/search/. **A.** The user selects two nodes. Here, the user is interested in Alzheimer’s disease, so [selects this](https://het.io/search/?source=17287) as the source node. The user limits the target node search to metanodes relating to gene function. The target node search box suggests nodes, sorted by the number of significant metapaths. When the user types in the target node box, the matches reorder based on search word similarity. Here, the user becomes interested in how the circadian rhythm might relate to Alzheimer’s disease. **B.** The webapp returns metapaths between Alzheimer’s disease and the circadian rhythm pathway. The user unchecks “precomputed only” to compute results for all metapaths with length ≤ 3, not just those that surpass the database inclusion threshold. The user sorts by adjusted *p*-value and selects 7 of the top 10 metapaths. **C.** Paths for the selected metapaths are ordered by their path score. The user selects 8 paths (1 from a subsequent page of results) to show in the graph visualization and highlights a single path involving *ARNT2* for emphasis. **D.** A subgraph displays the previously selected paths. The user improves on the automated layout by repositioning nodes. Clicking an edge displays its properties, informing the user that association between Creutzfeldt-Jakob disease and *NPAS2* was detected by GWAS.

Figure 2 shows the information used to compute *p*-value for enriched metapaths. The table includes the following columns:

- **path count**: The number of paths between the source and target node of the specified metapath
- **adjusted *p*-value**: A measure of the significance of the DWPC that indicates whether more paths were observed than expected due to random chance. Compares the DWPC to a null distribution of DWPCs generated from degree-preserving permuted networks. Bonferroni-adjusted for the number of metapaths with the same source metanode, target metanode, and length.
- ***p*-value**: A measure of the significance of the DWPC that indicates whether more paths were observed than expected due to random chance. Compares the DWPC to a null distribution of DWPCs generated from degree-preserving permuted networks. Not adjusted for multiple comparisons (i.e. when multiple metapaths are assessed for significant connectivity between the source and target node).
- **DWPC**: Degree-Weighted Path Count — Measures the extent of connectivity between the source and target node for the given metapath. Like the path count, but with less weight given to paths along high-degree nodes.
- **source degree**: The number of edges from the source node that are of the same type as the initial metaedge of the metapath.
- **target degree**: The number of edges from the target node that are of the same type as the final metaedge of the metapath.
- # **DWPCs**: The number of DWPCs calculated on permuted networks used to generate a null distribution for the DWPC from the real network. Permuted DWPCs are aggregated for all permuted node pairs with the same degrees as the source and target node.
- # **non-0 DWPCs**: The number of permuted DWPCs from ‘# of DWPCs’ column that were nonzero. Nonzero DWPCs indicate at least one path between the source and target node existed in the permuted network.
- **non-0 mean**: The mean of nonzero permuted DWPCs. Used to generate the gamma-hurdle model of the null DWPC distribution.
- **non-0 σ**: The standard deviation of nonzero permuted DWPCs. Used to generate the gamma-hurdle model of the null DWPC distribution.
- **Neo4j Actions**: A Cypher query that users can run in the Neo4j browser to show paths with the largest DWPCs for the metapath.

### Enriched paths

In addition to knowing which metapaths are enriched between two query nodes, it is helpful to see the specific paths that contribute highly to such enrichment. Since the DWPC is a summation of a path metric (called the path degree product), it is straightforward to calculate the proportion of a DWPC attributable to an individual path. The webapp allows users to select a metapath to populate a table of the corresponding paths. These paths are generated on-the-fly through a Cypher query to the Hetionet Neo4j database.

It is desirable to have a consolidated view of paths across multiple metapaths. Therefore, we calculate a *path score* heuristic, which can be used to compare the importance of paths between metapaths. The path score equals the proportion of the DWPC contributed by a path multiplied by the magnitude of the DWPC’s *p*-value (-log_10_(*p*)). To illustrate, the paths webapp panel includes the following information (Figure 4C):

- **path**: The sequence of edges in the network connecting the source node to the target node. Duplicate nodes are not permitted in paths.
- **path score**: A metric of how meaningful the path is in describing the connectivity between the source and target node. The score combines the magnitude of the metapath’s p-value with the percent of the DWPC contributed by the path.
- **% of DWPC** The contribution of the path to the DWPC for its metapath. This metric compares the importance of all paths of the same metapath from the source node to the target node.

### Hetio Website and Connectivity Search Webapp

We revamped the website hosted at https://het.io to serve as a unified home for this study and the hetnet-related research that preceded it. The website provides the connectivity search webapp running over the hetio network and several other interactive apps for prior projects. It also includes high-level information on hetnets and Hetionet, citation and contact details, links to supporting studies and software, downloads and exploration of Hetionet data, and related media.

We created the connectivity search webapp available at https://het.io/search/. The tool is free to use, without any login or authentication. The app allows users to quickly explore how any two nodes in Hetionet v1.0 might be related. The workflow accepts one or more nodes as input and shows the user the most important metapaths and paths for a pair of query nodes.

The design guides the user through selecting a source and target node (Figure 4A). The webapp returns metapaths, scored by whether they occurred more than expected based on network degree (Figure 4B). Users can proceed by requesting the specific paths for each metapath, which are placed in a unified table sorted according to their path score (Figure 4C). Finally, the webapp produces publication-ready visualizations containing user-selected paths (Figure 4D).

## Discussion

In this study, we introduce a search engine for hetnet connectivity between two nodes that returns results in realtime. An interactive webapp helps users explore node connectivity by ranking metapaths and paths, while visualizing multiple paths in a subgraph.

We made several methodological contributions to support this application. We developed optimized algorithms for computing DWPCs using matrix multiplication. In addition, we created a method for estimating a *p*-value for a DWPC, using null DWPCs computed on permuted hetnets. We implemented these advances in the open-source hetmatpy Python package and HetMat data structure to provide highly-optimized computational infrastructure for representing and reasoning on hetnets using matrices.

This work lays the foundation for exciting future directions. Here, we computed all DWPCs for Hetionet metapaths with length ≤ 3. Our search engine will therefore overlook important connectivity from longer metapaths. However, it is infeasible to compute DWPCs for all longer metapaths. One solution would be to only extend metapaths detected as informative. For example, if a *CbGpPWpG* metapath is deemed informative, it could be extended with additional metaedges like *CbGpPWpGaD*. One unsupervised approach would be to use the distribution of DWPC *p*-values for a metapath to detect whether the paths still convey sufficient information, for example by requiring an enrichment of small *p*-values. Were this method to fail, supervised alternatives could be explored, such as the ability for DWPCs from a longer metapath to predict that of a shorter metapath or metaedge, with care taken to prevent label leakage. One final approach could learn from user interest and compute longer metapaths only when requested.

This work focuses on queries where the input is a node pair. Equally interesting would be queries where the input is a set of nodes of the same type, optionally with weights. The search would compute DWPCs for paths originating on the query nodes. The simpler formulation would compute DWPCs for metapaths separately and compare to null distributions from permuted hetnets. A more advanced formulation would combine scores across metapaths such that every node in the hetnet would receive a single score capturing its connectivity to the query set. This approach would have similar utility to gene set enrichment analysis (GSEA) in that the user could provide a set of genes as input and receive a ranked list of nodes that characterize the function of the query genes. However, it would excel in its versatility by returning results of any node type without requiring pre-defined gene sets to match against. Some users might be interested in node set transformations where scores for one node type are converted to another node type. This approach could take scores for human genes and convert them to side effects, diseases, pathways, etcetera.

Our work is not without limitations. The final application relies on multiple databases and cached computations specific to Hetionet v1.0. Despite striving for a modular architecture, generating an equivalent search webapp for a different hetnet would require adaptation due to the many data sources involved. Furthermore, we would benefit from greater real-world evaluation of the connectivity search results to help identify situations where the method underperforms. Despite these challenges, our study demonstrates one of the first public search engines for node connectivity on a biomedical knowledge graph, while contributing methods and software that we hope will inspire future work.

## Methods

### Hetionet

We used the hetionet knowledge graph to demonstrate connectivity search. [Hetionet](https://het.io/about/) is a knowledge graph of human biology, disease, and medicine, integrating information from millions of studies and decades of research. Hetionet v1.0 combines information from [29 public databases](https://git.dhimmel.com/rephetio-manuscript/#tbl:resources). The network contains 47,031 nodes of [11 types](https://git.dhimmel.com/rephetio-manuscript/#tbl:metanodes) (Table 1) and 2,250,197 edges of [24 types](https://git.dhimmel.com/rephetio-manuscript/#tbl:metaedges) (Figure 1A).

**Table 1:** Node types in Hetionet. The abbreviation, number of nodes, and description for each of the 11 metanodes in Hetionet v1.0.

| Metanode | Abbr | Nodes | Description |
| --- | --- | --- | --- |
| Anatomy | A | 402 | Anatomical structures, excluding structures that are known not to be found in humans. From Uberon. |
| Biological Process | BP | 11381 | Larger processes or biological programs accomplished by multiple molecular activities. From Gene Ontology. |
| Cellular Component | CC | 1391 | The locations relative to cellular structures in which a gene product performs a function. From Gene Ontology. |
| Compound | C | 1552 | Approved small molecule compounds with documented chemical structures. From DrugBank. |
| Disease | D | 137 | Complex diseases, selected to be distinct and specific enough to be clinically relevant yet general enough to be well annotated. From Disease Ontology. |
| Gene | G | 20945 | Protein-coding human genes. From Entrez Gene. |
| Molecular Function | MF | 2884 | Activities that occur at the molecular level, such as “catalysis” or “transport”. From Gene Ontology. |
| Pathway | PW | 1822 | A series of actions among molecules in a cell that leads to a certain product or change in the cell. From WikiPathways, Reactome, and Pathway Interaction Database. |
| Pharmacologic Class | PC | 345 | “Chemical/Ingredient”, “Mechanism of Action”, and “Physiologic Effect” FDA class types. From DrugCentral. |
| Side Effect | SE | 5734 | Adverse drug reactions. From SIDER/UMLS. |
| Symptom | S | 438 | Signs and Symptoms (i.e. clinical abnormalities that can indicate a medical condition). From the MeSH ontology. |

One limitation that restricts the applicability of Hetionet is incompleteness. In many cases, Hetionet v1.0 includes only a subset of the nodes from a given resource. For example, the Disease Ontology contains over 9,000 diseases [31], while Hetionet includes only 137 diseases [32]. Nodes were excluded to avoid redundant or overly specific nodes, while ensuring a minimum level of connectivity for compounds and diseases. See the [Project Rephetio methods](https://git.dhimmel.com/rephetio-manuscript/#nodes) for more details [2]. Nonetheless, Hetionet v1.0 remains one of the most comprehensive and integrative networks that consolidates biomedical knowledge into a manageable number of node and edge types [33]. Other integrative resources, some still under development, include [Wikidata](https://www.wikidata.org/) [34], [SemMedDB](https://skr3.nlm.nih.gov/SemMedDB/) [35,36,37]. [SPOKE](https://spoke.ucsf.edu/), and [RTX-KG2c](https://github.com/RTXteam/RTX-KG2) [38].

### HetMat architecture

At the core of the hetmatpy package is the HetMat data structure for storing and accessing the network. HetMats are stored on disk as a directory, which by convention uses a `.hetmat` extension. A HetMat directory stores a single heterogeneous network, whose data resides in the following files.

1. A `metagraph.json` file stores the schema, defining which types of nodes and edges comprise the hetnet. This format is defined by the hetnetpy Python package. Hetnetpy was originally developed with the name hetio during prior studies [2,39], but we renamed it to het**net**py for better disambiguation from het**mat**py.
2. A `nodes` directory containing one file per node type (metanode) that defines each node. Currently, `.tsv` files where each row represents a node are supported.
3. An `edges` directory containing one file per edge type (metadata) that encodes the adjacency matrix. The matrix can be serialized using either the Numpy dense format (`.npy`) or SciPy sparse format (`.sparse.npz`).

For node and edge files, compression is supported as detected from `.gz`, `.bz2`, `.zip`, and `.xz` extensions. This structure of storing a hetnet supports selectively reading nodes and edges into memory. For example, a certain computation may only require access to a subset of the node and edge types. By only loading the required node and edge types, we reduce memory usage and read times.

Additional subdirectories, such as `path-counts` and `permutations`, store data generated from the HetMat. By using consistent paths for generated data, we avoid recomputing data that already exists on disk. A HetMat directory can be zipped for archiving and transfer. Users can selectively include generated data in archives. Since the primary application of HetMats is to generate computationally demanding measurements on hetnets, the ability to share HetMats with precomputed data is paramount.

The `HetMat` class implements the above logic. A `hetmat_from_graph` function creates a HetMat object and directory on disk from the pre-existing `hetnetpy.hetnet.Graph` format.

We converted Hetionet v1.0 to HetMat format and uploaded the `hetionet-v1.0. hetmat.zip` archive to the [Hetionet data repository](https://github.com/hetio/hetionet/tree/master/hetnet/matrix).

### DWPC matrix multiplication algorithms

Prior to this study, we used two implementations for computing DWPCs. The first is a pure Python implementation available in the `hetnetpy.pathtools.DWPC` function [39]. The second uses a Cypher query, prepared by `hetnetpy.neo4j.construct_dwpc_query`, that is executed by the Neo4j database [2,40]. Both of these implementations require traversing all paths between the source and target node. Hence, they are computationally cumbersome despite optimizations [41].

Since our methods only require degree-weighted counts, not fully enumerated paths, adjacency matrix multiplication presents an alternative approach. Multiplication alone, however, counts walks rather than paths, meaning paths traversing a single node multiple times are counted. When computing network-based features to quantify the relationship between a source and target node, we would like to exclude traversing duplicate nodes (i.e. paths, not trails nor walks) [42]. We developed a suite of algorithms to compute true path counts and DWPCs using matrix multiplication that benefits from the speed advantages of only counting paths.

Our implementation begins by categorizing a metapath according to the pattern of its repeated metanodes, allowing DWPC computation using a specialized order of operations. For example, the metapath *DrDtCrC* is categorized as a set of disjoint repeats, while *DtCtDpC* is categorized as repeats of the form BABA. Many complex repeat patterns can be represented piecewise as simpler patterns, allowing us to compute DWPC for most metapaths up to length 5 and many of length 6 and beyond without enumerating individual paths. For example, disjoint groups of repeats like *DrDtCrC* can be computed as the matrix product of DWPC matrices for *DrD* and *CrC*. Randomly-inserted non-repeated metanodes (e.g. *G* in *DrDaGaDrD)* require no special treatment and are included in DWPC with matrix multiplication.

After metapath categorization, we segment metapaths according to their repeat pattern, following our order of operations. By segmenting and computing recursively, we can evaluate DWPC efficiently on highly complex metapaths, using simple patterns as building-blocks for higher-level patterns. Finally, our specialized DWPC functions are applied to individual segments, the results are combined, and final corrections are made to ensure no repeated nodes are counted. The recursive, segmented approach we developed allowed us additionally to implement a caching strategy that improved speed by avoiding duplicate DWPC computations. In summary, the functionality we developed resulted in greater than a 175-fold reduction in compute time, allowing us to compute millions of DWPC values across Hetionet [43].

### Details of matrix DWPC implementation

DWPC computation requires us to remove all duplicate nodes from paths. We used three repeat patterns as the building blocks for DWPC computation: short repeats (AAA), nested repeats (BAAB), and overlapping repeats (BABA). Let **• •*XwXyX*•** denote the DWPC matrix for metapath *XwXyZ*. Under this notation, **• •(*XyZ*)•** is the degree-weighted (bi)adjacency matrix for metaedge *XyZ*. Additionally, let **• •• • •*A*•** represent a diagonal matrix whose entries are the diagonal elements of *A*.

For the case of short (< 4) repeats for a single metanode, *XaXbX*(e.g. *GiGdG)*, we simply subtract the main diagonal.

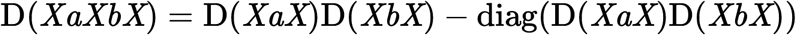

Nested repeats *XaYbYcX*(e.g. *CtDrDtC*), are treated recursively, with both inner (YY) and outer (XX) repeats treated as separate short repeats.

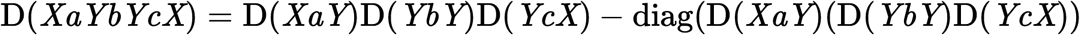

Overlapping repeats *XaYbXcY*(e.g. *CtDtCtD*) require several corrections (• denotes the Hadamard product).

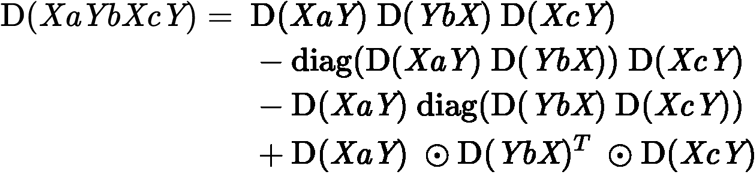

Most paths of length six—and many even longer paths—can be represented hierarchically using these patterns. For example, a long metapath pattern of the form CBABACXYZ can be segmented as (C(BABA)C)XYZ using patterns for short and overlapping repeats and can be computed using the tools we developed. In addition to these matrix routines—which advantageously count rather than enumerate paths—we implemented a general matrix method for any metapath type. The general method is important for patterns such as long (≥ 4) repeats, or complex repeat patterns (e.g. of the form ABCABC), but it requires path enumeration and is therefore slower. As an alternative approach for complex paths, we developed an approximate DWPC method that corrects repeats in disjoint simple patterns but only corrects the first repeat in complex patterns (e.g. ≥ length four repeat). Mayers et al. developed an alternative approximation, which subtracts the main diagonal at every occurrence of the first repeated metanode [44]. All our matrix methods were validated against existing implementations involving explicit path enumeration to ensure consistent results.

### Permuted hetnets

In order to generate a null distribution for a DWPC, we rely on DWPCs computed from permuted hetnets. We derive permuted hetnets from the unpermuted network using the XSwap algorithm [45]. XSwap randomizes edges while preserving node degree. Therefore, it’s ideal for generating null distributions that retain general degree effects, but destroy the actual meaning of edges. We adapt XSwap to hetnets by applying it separately to each metaedge [2,46,47].

Project Rephetio created 5 permuted hetnets [2,46], which were used to generate a null distribution of classifier performance for each metapath-based feature. Here, we aim to create a null distribution for individual DWPCs, which requires vastly more permuted values to estimate with accuracy. Therefore, we generated 200 permuted hetnets (archive). More recently, we also developed the `xswap` Python [package](https://github.com/greenelab/xswap), whose optimized C/C++ implementation will enable future research to generate even larger sets of permuted networks [47].

### Degree-grouping of node pairs

For each of the 200 permuted networks and each of the 2,205 metapaths, we computed the entire DWPC matrix (i.e. all source nodes × target nodes). Therefore, for each actual DWPC value, we computed 200 permuted DWPC values. Because permutation preserves only node degree, DWPC values among nodes with the same source and target degrees are equivalent to additional permutations. We greatly increased the effective number of permutations by grouping DWPC values according to node degree, affording us a superior estimation of the DWPC null distribution.

We have applied this *degree-grouping* approach previously when calculating the prior probability of edge existence based on the source and target node degrees [47,48]. But here, we apply *degree-grouping* to null DWPCs. The result is that the null distribution for a DWPC is based not only on permuted DWPCs for the corresponding source–metapath–target combination, but instead on all permuted DWPCs for the source-degree–metapath–target-degree combination.

The “# DWPCs” column in Figure 2 illustrates how degree-grouping inflates the sample size of null DWPCs. The *p*-value for the *DaGiGpPW* metapath relies on the minimum number of null DWPCs (200), since no other disease besides Alzheimer’s had 196 *associates* edges (source degree) and no other pathway besides circadian rhythm had 201 *participates* edges (target degree). However, for other metapaths with over 5,000 null DWPCs, degree-grouping increased the size of the null distribution by a factor of 25. In general, source–target node pairs with lower degrees receive the largest sample size multiplier from degree-grouping. This is convenient since low-degree nodes also tend to produce the highest proportion of zero DWPCs, by virtue of low connectivity. Consequently, degree grouping excels where it is needed most.

One final benefit of degree-grouping is that reduces the disk space required to store null DWPC summary statistics. For example, with 20,945 genes in Hetionet v1.0, there exists 438,693,025 gene pairs. Gene nodes have 302 distinct degrees for *interacts* edges, resulting in 91,204 degree pairs. This equates to an 4,810-fold reduction in the number of summary statistics that need to be stored to represent the null DWPC distribution for a metapath starting and ending with a *Gene-interacts-Gene* metaedge.

We store the following null DWPC summary statistics for each metapath-source-degree-target-degree combination: total number of null DWPCs, total number of nonzero null DWPCs, sum of null DWPCs, sum of squared null DWPCs, and number of permuted hetnets. These values are sufficient to estimate the *p*-value for a DWPC, by either [fitting](https://github.com/hetio/hetmatpy/blob/bc36aa9859c43a1a5fb22808cd6eb952ef9d497c/hetmatpy/pipeline.py#L42-L63) a gamma-hurdle null distribution or generating an empiric *p*-value. Furthermore, these statistics are additive across permuted hetnets. Their values are always a running total and can be updated incrementally as statistics from each additional permuted hetnet become available.

Figure 5 shows how various aspects of DWPCs vary by degree group. The rows display the following metrics of the DWPC distribution for all node-pairs in a given degree-group:

- # **Nonzero DWPCs**: The number of nonzero DWPCs values (on average per network to enable comparison).
- **% Nonzero DWPCs**: Of the total number of DWPCs, the percent that is nonzero. As node degrees increase, the chance of node pairs having at least one path, and hence a nonzero DWPC, greatly increases.
- **Mean DWPC**: The average value of all DWPCs including zeros.
- **Mean Nonzero DWPC**: The average value of nonzero DWPCs.
- **Std Dev Nonzero DWPC**: The standard deviation of nonzero DWPCs.
- **Gamma Model β**: The β parameter of the gamma model fìt on nonzero DWPCs. Note that the gamma model is only fìt on permuted network DWPCs to estimate a null distribution for the unpermuted network DWPCs. Since this parameter varies with source & target node degree, it is important to fìt a separate gamma model for each degree group.

**Figure 5:**
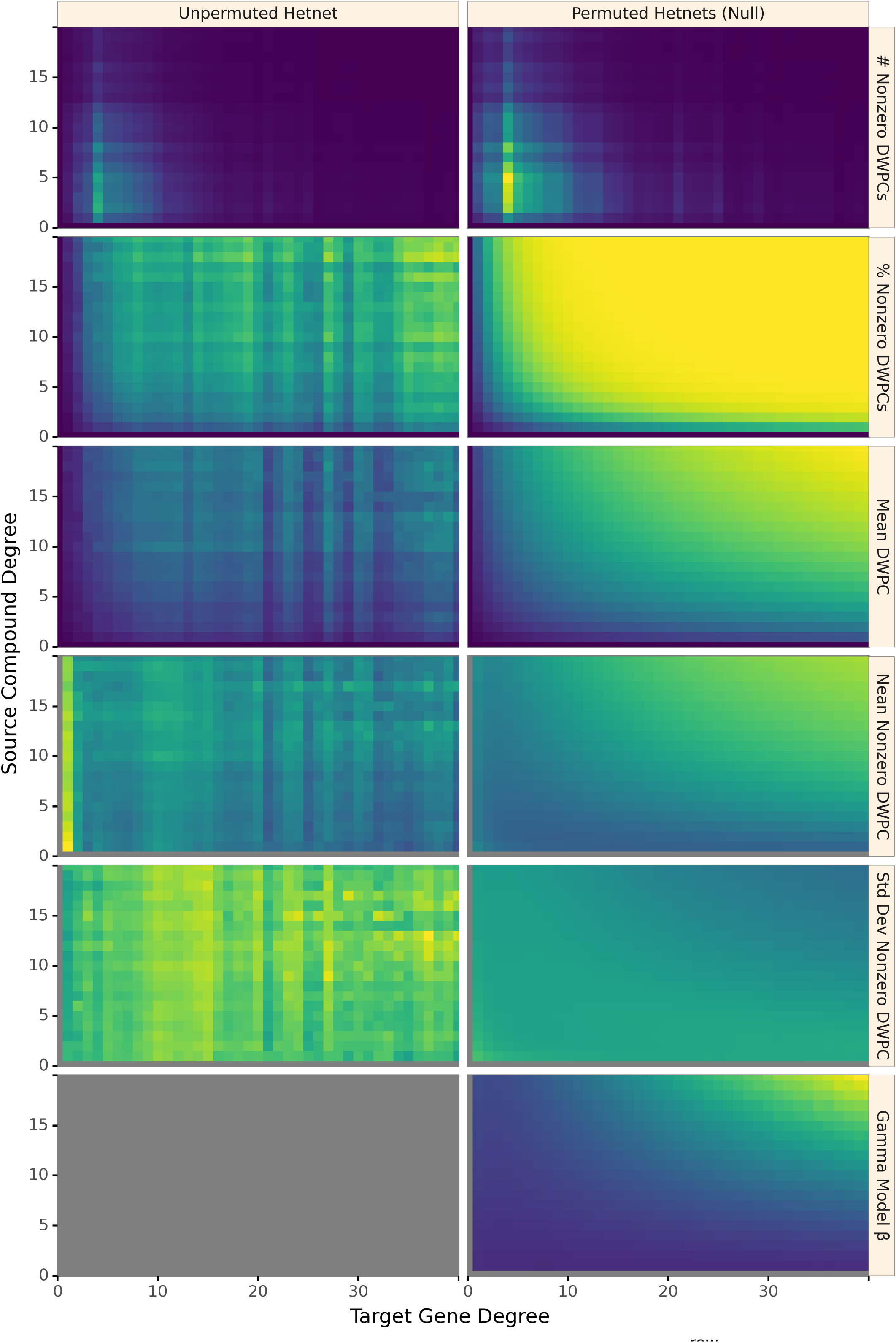
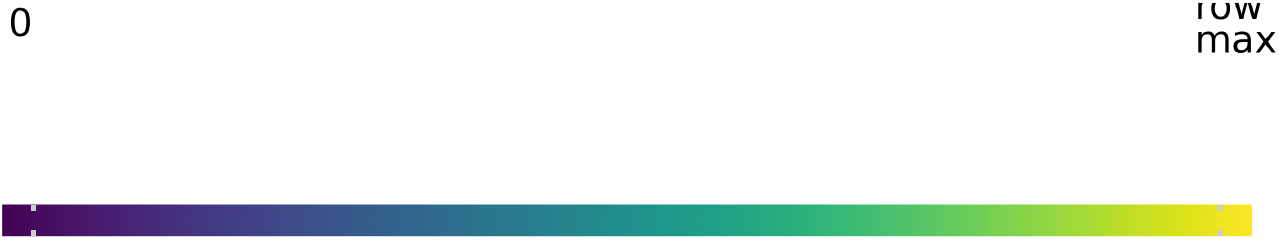
Path-based metrics vary by node degree and network permutation status. Each row shows a different metric of the DWPC distribution for the *CbGpPWpG* metapath — traversing Compound–binds–Gene–participates–Pathway–participates–Gene, selected for illustrative purposes. Metrics are computed for degree-groups, which is a specific pair of source degree (in this case, the source compound’s count of CbG edges) and target degree (in this case, the target gene’s count of GpPW edges). On the left, metrics are reported for the unpermuted hetnet and on the right for the 200 permuted hetnets. Hence, each cell on right summarizes 200 times the number of DWPCs as the corresponding cell on the left. The colormap is row normalized, such that its intensity peaks for the maximum value of each metric across the unpermuted and permuted values. Gray indicates null values.

### Gamma-hurdle distribution

We are interested in identifying source and target nodes whose connectivity exceeds what typically arises at random. To identify such especially-connected nodes, we compare DWPC values to the distribution of permuted network DWPC values for the same source and target nodes. While a single DWPC value is not actually a test statistic, we use a framework akin to classical hypothesis testing to identify outliers.

Two observations led us to the quasi-significance testing framework we developed. First, a sizable fraction of permuted DWPC values is often zero, indicating that the source and target nodes are not connected along the metapath in the permuted network. Second, we observed that non-zero DWPC values for any given source and target nodes are reasonably approximated as following a gamma distribution. Motivated by these observations, we parametrized permuted DWPC values using a zero-inflated gamma distribution, which we termed the gamma-hurdle distribution. We fit a gamma-hurdle distribution to each combination of source node, target node, and metapath. Finally, we estimate the probability of observing a permuted DWPC value greater than DWPC computed in the unpermuted network, akin to a one-tailed p-value. These quasi-significance scores (‘*p*-values’) allow us to identify outlier node pairs at the metapath level (see examples in Figure 6).

**Figure 6:**
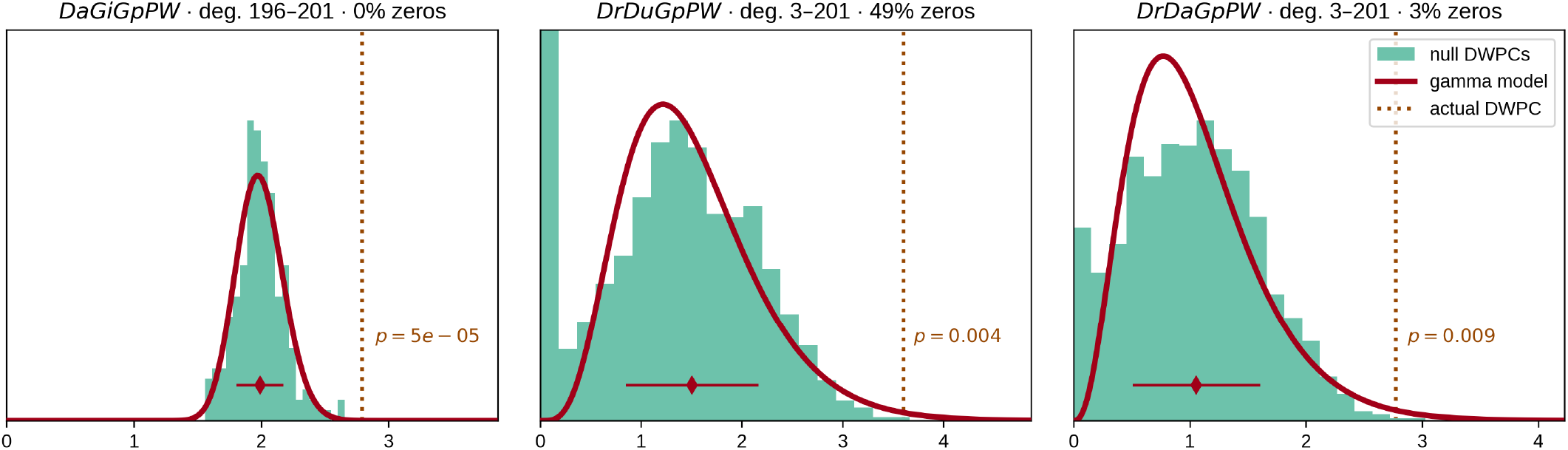
From null distribution to *p*-value for DWPCs. Null DWPC distributions are shown for three metapaths between Alzheimer’s disease and the circadian rhythm pathway, selected from Figure 2. For each metapath, null DWPCs are computed on 200 permuted hetnets and grouped according to source–target degree. Histograms show the null DWPCs for the degree group corresponding to Alzheimer’s disease and the circadian rhythm pathway (as noted in the plot titles by deg.) The proportion of null DWPCs that were zero is calculated, forming the “hurdle” of the null distribution model. The nonzero null DWPCs are modeled using a gamma distribution, which can be fit solely from a sample mean and standard deviation. The mean of nonzero null DWPCs is denoted with a diamond, with the standard deviation plotted twice as a line in either direction. Actual DWPCs are compared to the gamma-hurdle null distribution to yield a *p*-value.

### Details of the gamma-hurdle distribution

Let *X*be a gamma-hurdle random variable with parameters *λ, α*, and *β*.

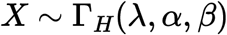

The probability of a draw from the distribution is

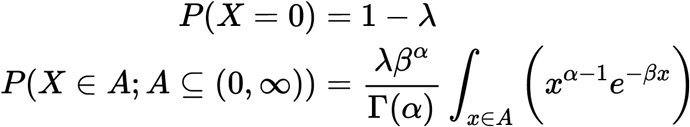

We estimate all three parameters using the method of moments (using Bessel’s correction to estimate the second moment). As a validation of our method, we [compared](https://nbviewer.jupyter.org/github/greenelab/hetmech/blob/025bdf8d5e63725ca2482d61fd8e421bf0001f93/explore/gamma-hurdle/parameter_estimates.ipynb) our method of moments parameter estimates to approximate maximum likelihood estimates (gamma distribution parameters do not have closed-form maximum likelihood estimates) and found excellent concordance between the methods. Let *N* be the number of permuted DWPC values, and *n* the number of nonzero values.

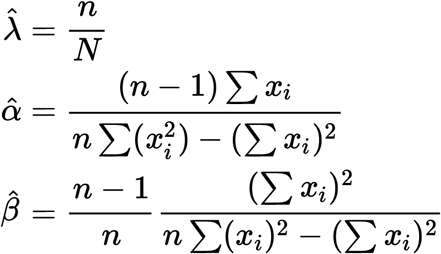

Finally, we compute a p-value for each DWPC value, *t*.

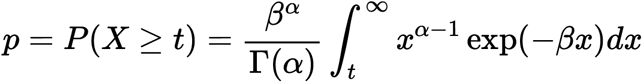

### Empirical DWPC p-values

We [calculate](https://github.com/hetio/hetmatpy/blob/bc36aa9859c43a1a5fb22808cd6eb952ef9d497c/hetmatpy/pipeline.py#L92-L113) an empirical p-value for special cases where the gamma-hurdle model cannot be applied. These cases include when the observed DWPC is zero or when the null DWPC distribution is all zeroes or has only a single distinct nonzero value. The empirical *p*-value (*p_empiric_*) equals the proportion of null DPWCs ≥ the observed DWPC.

Since we don’t store all null DWPC values, we apply the following criteria to calculate *p_empiric_* from summary statistics:

1. When the observed DWPC = 0 (no paths of the specified metapath existed between the source and target node), *p_empiric_* = 1.
2. When all null DWPCs are zero but the observed DWPC is positive, *p_empiric_* = 0.
3. When all nonzero null DWPCs have the same positive value (standard deviation = 0), *p_empiric_* = 0 if the observed DWPC > the null DWPC else *p_empiric_* = proportion of nonzero null DWPCs.

### DWPC and null distribution computation

We decided to compute DWPCs and their significance for all source–target node pairs for metapaths with length **≤** 3. On Hetionet v1.0, there are 24 metapaths of length 1, 242 metapaths of length 2, and 1,939 metapaths of length 3. The decision to stop at length 3 was one of practicality, as length 4 would have added 17,511 metapaths.

For each of the 2,205 [metapaths](https://github.com/greenelab/hetmech/raw/042063fb559048e52b3dc2731b6d6c6836f698cf/explore/bulk-pipeline/archives/metapath-dwpc-stats.tsv), we computed the complete path count matrix and DWPC matrix ([notebook](https://nbviewer.jupyter.org/github/greenelab/hetmech/blob/042063fb559048e52b3dc2731b6d6c6836f698cf/explore/bulk-pipeline/bulk.ipynb)). In total, we computed 137,786,767,964 path counts (and the same number of DWPCs) on the unpermuted network, of which 11.6% were nonzero.

The DWPC has a single parameter, called the damping exponent (*w*), which controls how much paths through high-degree nodes are downweighted [39]. When *w* = 0, the DWPC is equivalent to the path count. Previously, we found *w* = 0.4 was optimal for predicting disease-associated genes [39]. Here, we use *w* = 0.5, since taking the square root of degrees has more intuitive appeal.

We selected data types for matrix values that would allow for high precision. We used 64-bit unsigned integers for path counts and 64-bit floating-point numbers for DWPCs. We [considered](https://github.com/greenelab/hetmech/pull/91) using 16-bits or 32-bits per DWPC to reduce memory/storage requirements, but decided against it in case certain applications required greater precision.

We used SciPy sparse for path count and DWPC matrices with density < 0.7, serialized to disk with compression and a `.sparse.npz` extension. This format minimizes the space on disk and load time for the entire matrix but does not offer read access to slices. We used Numpy 2D arrays for DWPC matrices with density ≥ 0.7, serialized to disk using Numpy’s `.npy` format. We bundled the path count and DWPC matrix files into HetMat archives by metapath length and deposited the archives to Zenodo [49]. The archive for length 3 DWPCs was the largest at 131.7 GB.

We also generated null DWPC summary statistics for the 2,205 metapaths, which are also available by metapath length from Zenodo as HetMat archives consisting of `.tsv.gz` files [49]. Due to degree grouping, null DWPCs summary statistic archives are much smaller than the DWPC archives. The archive for length 3 null DWPCs summary statistics was 733.1 MB. However, the compute required to generate null DWPCs is far greater because there are multiple permuted hetnets (in our case 200). As a result, computing and saving all DWPCs took 6 hours, whereas computing and saving the null DWPC summary statistics took 361 hours.

Including null DWPCs and path counts, the Zenodo deposit totals 185.1 GB and contains the results of computing ~28 trillion DWPCs — 27,832,927,128,728 to be exact.

### Adjusting DWPC *p*-values

When a user applies hetnet connectivity search to identify enriched metapaths between two nodes, many metapaths are evaluated for significance. Due to multiple testing of many DWPCs, low *p*-values are likely to arise by chance. Therefore, we devised a multiple-testing correction.

For each combination of source metanode, target metanode, and length, we counted the number of metapaths. For Disease…Pathway metapaths, there are 0 metapaths of length 1, 3 metapaths of length 2, and 24 metapaths of length 3. We calculated adjusted p-values by applying a Bonferroni correction based on the number of metapaths of the same length between the source and target metanode. Using Figure 2 as an example, the DdGpPW p-value of 5.9% was adjusted to 17.8% (multiplied by a factor of 3).

Bonferroni controls family-wise error rate, which corresponds here to incorrectly finding that *any* metapath of a given length is enriched. As a result, our adjusted p-values are conservative. We would prefer to adjust *p*-values for false discovery rate [50], but these methods often require access to all *p*-values at once (impractical here) and assume a uniform distribution of *p*-values when there is no signal (not the case here when most DWPCs are zero).

### Prioritizing enriched metapaths for database storage

Storing DWPCs and their significance in the database (as part of the `PathCount` table in Figure 7) enables the connectivity search webapp to provide users with enriched metapaths between query nodes in real time. However, storing ~15.9 billion rows (the total number of nonzero DWPCs) in the database’s `PathCount` table would exceed a reasonable disk quota. An alternative would be to store all DWPCs in the database whose adjusted p-value exceeded a universal threshold (e.g. *p* < 5%). But we estimated this would still be prohibitively expensive. Therefore, we devised a metapath-specific threshold. For metapaths with length 1, we stored all nonzero DWPCs, assuming users always want to be informed about direct edges between the query nodes, regardless of significance. For metapaths with length ≥ 2, we chose an adjusted *p*-value threshold of 5 × (*n_source_* × *n_target_*)^-0.3^, where *n_source_* and *n_target_* are the node counts for the source and target metanodes (i.e. “Nodes” column in Table 1). Notice that metapaths with large number of possible source–target pairs (large DWPC matrices) are penalized. This decision is based on practicality since otherwise the majority of the database quota would be consumed by a minority of metapaths between plentiful metanodes (e.g. Gene…Gene metapaths). Also, we assume that users will search nodes at a similar rate by metanode (e.g. they’re more likely to search for a specific disease than a specific gene). The constants in the threshold formula help scale it. The multiplier of 5 relaxes the threshold to saturate the available database capacity. The −0.3 exponent applies the large DWPC-matrix penalty.

**Figure 7:**
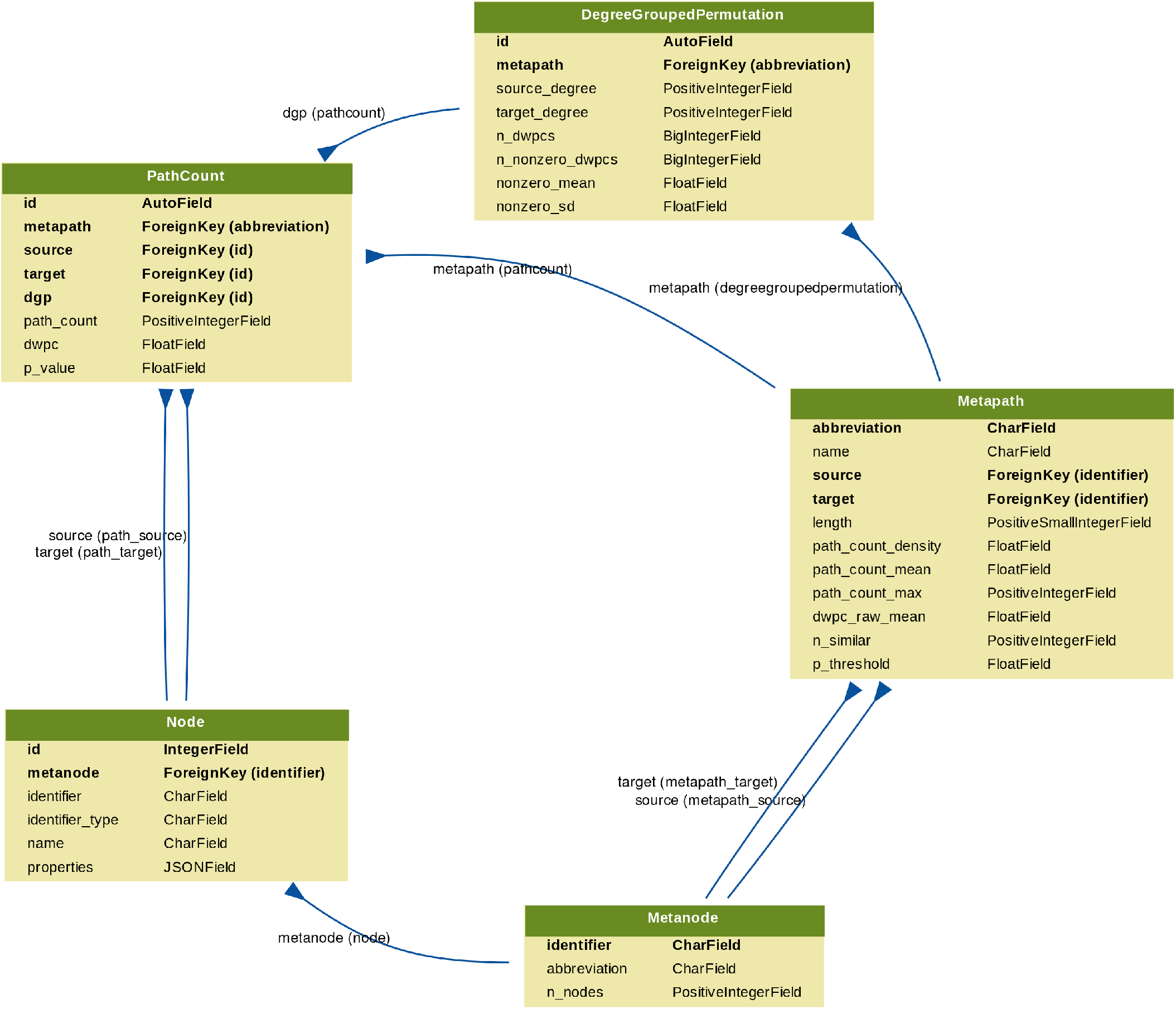
Schema for the connectivity search backend relational database models. Each Django model is represented as a table, whose rows list the model’s field names and types. Each model corresponds to a database table. Arrows denote foreign key relationships. The arrow labels indicate the foreign key field name followed by reverse relation names generated by Django (in parentheses).

Users can still evaluate DWPCs that are not stored in the database, using either the webapp or API. These are calculated on the fly, delegating DWPC computation to the Neo4j database. Unchecking “precomputed only” on the webapp shows all possible metapaths for two query nodes. For some node pairs, the on-the-fly computation is quick (less than a second). Other times, computing DWPCs for all metapaths might take more than a minute.

### Backend Database & API

We created a backend application using Python’s Django web framework. The source code is available in the connectivity-search-backend repository. The primary role of the backend is to manage a relational database and provide an API for requesting data.

We define the database schema [using](https://github.com/greenelab/connectivity-search-backend/blob/af12f8cf2ad47d9a25ce8d1b7889390654eb3bb9/dj_hetmech_app/models.py) Django’s object-relational mapping framework (Figure 7). We [import](https://github.com/greenelab/connectivity-search-backend/blob/af12f8cf2ad47d9a25ce8d1b7889390654eb3bb9/dj_hetmech_app/management/commands/populate_database.py) the data into a PostgreSQL database. Populating the database for all 2,205 metapaths up to length 3 was a prolonged operation, [taking](https://github.com/greenelab/connectivity-search-backend/pull/41#issuecomment-488054789) over 3 days. The majority of the time is spent populating the `DegreeGroupedPermutation` (37,905,389 rows) and `PathCount` (174,986,768 rows) tables. To avoid redundancy, the database only stores a single orientation of a metapath. For example, if rows are stored for the *GpPWpGaD* metapath, they would not also be stored for the *DaGpPWpG* metapath. The backend is responsible for checking both orientations of a metapath in the database and reversing metapaths on-the-fly before returning results. The database is located at `search-db.het.io` with public read-only access [available](https://github.com/greenelab/connectivity-search-backend#database).

We host a public API instance at https://search-api.het.io. Version 1 of the API exposes several endpoints that are used by the connectivity search frontend including queries for node details (/v1/node), node lookup (/v1/nodes), metapath information (/v1/metapaths), and path information (/v1/paths). The endpoints return JSON payloads. Producing results for these queries relies on internal calls to the PostgreSQL relational database as well as the pre-existing Hetionet v1.0 Neo4j graph database. They were designed to power the hetnet connectivity search webapp, but are also available for general research use.

### Frontend

#### Hetio Website

We created a static website to serve as the home for the Hetio organization using Jekyll. The source code is available in the het.io repository.

To ease the burden of maintenance and typical website hosting costs, the HTML, CSS, JavaScript, and other assets for the website are hosted for free on GitHub Pages. Jekyll was chosen over other static site generators for simplicity, ease of use, popularity (support), and its convenient integration with GitHub Pages. To make a change to the website, an author simply commits the changes (either directly or through a pull request) to the repository’s `gh-pages` branch, and GitHub automatically recompiles the website and hosts the resulting files at the provided custom domain URL. No explicit build instructions or other continuous integration is required.

#### Webapps

We developed the connectivity search app as a single-page, standalone application using React and associated tools. The source code is available in the connectivity-search-frontend repository.

Since the rest of the overarching Hetio website was mostly non-interactive content, it was appropriate to construct the bulk of the website in simple static formats like Markdown and HTML using Jekyll, and leave React for implementing the sections of the site that required more complex behavior.

We used React’s own `create-react-app` command line tool to generate a boilerplate for the app. This greatly simplified setting up and maintaining the app’s testing and building pipelines, bypassing time-consuming configuration of things like Webpack and linters. Some configuration was necessary to produce non-hashed, consistently named output files like `index.js` that could be easily and reliably referenced by and embedded into the Hetio Jekyll website.

For authoring components, we used React’s traditional class syntax. At the time of authoring the app, React Hooks were still nascent, thus the simpler and less-verbose functional syntax was not viable.

While writing this application, we also elected to re-write the pre-existing [Rephetio](https://het.io/disease-genes/) and [disease-associated genes](https://het.io/disease-genes/) apps in the same manner. We created a custom package of React components and utility functions that could be shared across the multiple interactive apps on the website. The package is located at and can be installed from the frontend-components repository. The package consists of interface “components” (building blocks) like buttons, sortable/searchable/paginated tables, etc., as well as utility functions for formatting data, debugging, etcetera. Each of the interactive apps import this package to reduce code repetition and to enforce a consistent style and behavior across the website.

For managing state in the connectivity search app, we used the Redux library. In general, Redux is a well-accepted approach to managing complex state. To be more explicit, Redux was chosen over vanilla React or other state management libraries for a few reasons:

1. The state in this app was very “global”, meaning most of it was needed by a lot of different parts of the app. Redux provides a convenient global “store” of state that is easily accessible to any component, avoiding the “prop-drilling” phenomenon.
2. The structure of the state is nested and complex. Redux’s “reducer” approach makes it cleaner to modify this type of data.
3. Redux’s approach to immutable state that is updated by actions and pure functions makes the application easier to debug. It is easy to get a clear timeline of how and when the state changed, and what changed it.

To create the graph visualization at the bottom of the app, the D3 library was used. D3 was chosen over other many other library choices for flexibility and comprehensiveness of features. At the time of development, no other library could be found that satisfied several core requirements:

1. SVG implementation for high-resolution, publication-ready figures.
2. Force-directed layout for untangling nodes.
3. Pinnable nodes and other physics customizations.
4. Customizable node and edge drag/hover/select behavior.
5. Intuitive pan/zoom view that worked on desktop and mobile.
6. Node and edge appearances that were completely customizable for alignment, text wrapping, color, outlines, fonts, arrowheads, non-colliding coincident edges, etc.

### Visual Design

A limited palette of colors was chosen to represent the different types of nodes (metanodes) in the Hetionet knowledge graph. These colors are listed and programmatically accessible in the [hetionet](https://github.com/hetio/hetionet) repository under `/describe/styles.json`.

At the time of developing connectivity search, Hetionet already had an established palette of colors (from Project Rephetio). To avoid confusion, we were careful to keep the general hue of each metanode color the same for backwards compatibility, e.g. genes stayed generally blue, diseases stayed generally brown. In this way, this palette selection was more of a modernization/refresh. For cohesiveness, accessibility, and aesthetic appeal, we used the professionally-curated Material Design palette as a source for the specific color values.

The palette is now used in all Hetio-related applications and materials. This is not just to maintain a consistent look and feel across the Hetio organization, but to convey clear and precise meaning. For example, the colors used in the metagraph in Figure 1A are exactly the same colors, and thus represent the same types of entities, as in *any part of the connectivity search app* (Figure 4).

Colors in the palette are also used in the Hetio logo (seen in Figure 3) and other miscellaneous logos and iconography across the website, to establish an identifiable brand for the Hetio organization as a whole.

### Realtime open science

This study was conducted entirely in the open via public GitHub repositories. We used GitHub Issues for discussion, leaving a rich online history of the scholarly process. Furthermore, most additions to the analyses were performed by pull request, whereby a contributor proposes a set of changes. This provides an opportunity for other contributors to review changes before they are officially accepted. For example, in [greenelab/hetmech#156](https://github.com/greenelab/hetmech/pull/156) @zietzm proposed a notebook to visualize parameters for null DWPC distributions. After @zietzm addressed @dhimmel’s comments, the pull request was approved and merged into the project’s main branch.

The manuscript for this study was written using [Manubot](https://manubot.org/), which allows authors to collaboratively write manuscripts on GitHub [51]. The Manubot-rendered manuscript is available at https://greenelab.github.io/connectivity-search-manuscript/. We encourage readers with feedback or questions to comment publicly via [GitHub Issues](https://github.com/greenelab/connectivity-search-manuscript/issues).

### Software & data availability

#### Hetio

*Hetio* is a superset/collection of hetnet-related research, tools, and datasets that, most notably, includes the Hetionet project itself and the connectivity search tool that are the focus of this manuscript. Most of the Hetio resources and projects can be found under the [Hetio GitHub](https://github.com/hetio) organization, with others being available under the [Greene Lab GitHub organization](https://github.com/greenelab), one of the collaborating groups. Information about Hetio is also displayed and disseminated on the Hetio website, as noted in the <u>Hetio Website</u> section.

#### Hetnet Connectivity Search

This study primarily involves the following repositories:

- greenelab/connectivity-search-manuscript: Source code for this manuscript. Best place for general comments or questions. CC BY 4.0 License.
- greenelab/hetmech: The initial project repository that contains research notebooks, dataset generation code, and exploratory data analyses. The hetmatpy package was first developed as part of this repository until its [relocation](https://github.com/hetio/hetmatpy/issues/1) in November 2018. BSD 3-Clause License.
- greenelab/connectivity-search-backend: Source code for the connectivity search database and API. BSD 3-Clause License.
- greenelab/connectivity-search-frontend: Source code for the connectivity search webapp. BSD 3-Clause License.
- hetio/hetmatpy: Python package for matrix storage and operations on hetnets. Released on PyPI. BSD 2-Clause Plus Patent License.
- hetio/hetnetpy Preexisiting python package for representing hetnets. Dependency of hetmatpy. Released on [PyPI](https://pypi.org/project/hetnetpy/). Dual licensed under BSD 2-Clause Plus Patent License and CC0 1.0 (public domain dedication).
- hetio/hetionet. Preexisiting data repository for Hetionet, including the public Neo4j instance and HetMat archives. CC0 1.0 License.
- hetio/het.io. Preexisiting source code for the https://het.io/website. CC BY 4.0 License.

The hetmech and hetionet repositories contain datasets related to this study. Large datasets were compressed and tracked with [Git LFS](https://git-lfs.github.com/) (Large File Storage). GitHub LFS had a max file size of 2 GB. Datasets exceeding this size, along with other essential datasets, are available from Zenodo [49].

